# The lamellipodium is a myosin independent mechanosensor

**DOI:** 10.1101/186437

**Authors:** Patrick W. Oakes, Tamara C. Bidone, Yvonne Beckham, Austin V. Skeeters, Guillermina R. Ramirez-San Juan, Stephen P. Winter, Gregory A. Voth, Margaret L. Gardel

**Affiliations:** Institute for Biophysical Dynamics, University of Chicago, and Chicago, IL 60637; James Franck Institute, University of Chicago, and Chicago, IL 60637; Department of Physics, University of Chicago, and Chicago, IL 60637; Department of Chemistry, University of Chicago, Chicago, IL 60637; Department of Physics & Astronomy, University of Rochester, and Rochester, NY 14627; Department of Biology, University of Rochester, Rochester, NY 14627; Interdisciplinary Scientist Training Program, University of Chicago, Chicago, IL 60637

**Keywords:** Mechanosensing, focal adhesions, lamellipodium, catch-bonds

## Abstract

The ability of adherent cells to sense changes in the mechanical properties of their extracellular environments is critical to numerous aspects of their physiology. It has been well documented that cell attachment and spreading are sensitive to substrate stiffness. Here we demonstrate that this behavior is actually biphasic, with a transition that occurs around a Young’s modulus of ∼7 kPa. Furthermore, we demonstrate that, contrary to established assumptions, this property is independent of myosin II activity. Rather, we find that cell spreading on soft substrates is inhibited due to reduced nascent adhesion formation within the lamellipodium. Cells on soft substrates display normal leading edge protrusion activity, but these protrusions are not stabilized due to impaired adhesion assembly. Enhancing integrin-ECM affinity through addition of Mn^2+^ recovers nascent adhesion assembly and cell spreading on soft substrates. Using a computational model to simulate nascent adhesion assembly, we find that biophysical properties of the integrin-ECM bond are optimized to stabilize interactions above a threshold matrix stiffness that is consistent with the experimentally observations. Together these results suggest that myosin II-independent forces in the lamellipodium are responsible for mechanosensation by regulating new adhesion assembly, which in turn, directly controls cell spreading. This myosin II-independent mechanism of substrate stiffness sensing could potentially regulate a number of other stiffness sensitive processes.

**Significance Statement:** Cell physiology can be regulated by the mechanics of the extracellular environment. Here, we demonstrate that cell spreading is a mechanosensitive process regulated by weak forces generated at the cell periphery and independent of motor activity. We show that stiffness sensing depends on the kinetics of the initial adhesion bonds that are subjected to forces driven by protein polymerization. This work demonstrates how the binding kinetics of adhesion molecules are sensitively tuned to a range of forces that enable mechanosensation.

## Introduction

The ability of cells to sense mechanical forces and convert them into biochemical responses regulates a plethora of physiological functions (1-3). In particular, cells respond to changes in the stiffness of the extracellular matrix (ECM) by altering a number of adhesion dependent behaviors, including spreading (4-12), migration (4, 13, 14), proliferation (15), differentiation (16, 17), and metastasis (18, 19). Matrix mechanosensing is thought to be mediated by focal adhesions, hierarchical organelles comprised of ∼150 proteins that facilitate dynamic and force-sensitive interactions between the extracellular matrix and the actin cytoskeleton (20-22). How these dynamic organelles mediate environmental sensing in a variety of physiological contexts, however, is still largely unknown.

Previous efforts have focused primarily on myosin-II-mediated mechanisms for substrate stiffness sensing (23-28). Stresses generated by myosin motors on the actin cytoskeleton are transmitted to the ECM via focal adhesions. These stresses, coupled with the matrix rigidity, impact the deformation and binding affinity of proteins within the focal adhesion (29-32). Changes in the composition and kinetics of proteins within focal adhesions are thought to variably regulate force transmission from the actin cytoskeleton and the matrix (33-35), leading many to describe focal adhesions as molecular clutches. Initial adhesion formation, however, occurs in the leading edge of the lamellipodium and is a myosin-independent process (36, 37). These structures, known as nascent adhesions, are instead subject to forces that primarily originate from polymerization of actin filaments. The contribution of nascent adhesions to mechanisms of substrate stiffness sensing have not been thoroughly explored.

One of the best characterized metrics of environmental sensing by adherent cells is their ability to attach and spread on ligand-coated substrates. The extent of cell spreading is controlled by the density and spatial organization of matrix ligands (38-40) as well as the rigidity of the substrate to which these ligands are attached (4-12). In the limit of soft substrates with a Young’s Modulus less than 500 Pa, cell spreading is inhibited. As the substrate stiffness increases, the spread area increases and ultimately plateaus (8-12). While previous reports have differed on the exact range of relevant stiffness which regulates this behavior, likely due to variances in experimental approaches (41), cell spreading remains a robust metric to study substrate stiffness sensing.

Here we study the mechanism regulating substrate stiffness dependent cell spreading. We find that NIH 3T3 cell spreading is acutely impacted as the Young’s modulus of the substrate increases from 5 to 8 kPa. On substrates with a stiffness less than 5 kPa, cells spread poorly. Average cell spread area increases on substrates stiffer than 5 kPa, plateauing on substrates stiffer than 8 kPa. Above this threshold, cell spread area remains constant. Surprisingly, we find this stiffness-dependent changes in cell spreading is independent of myosin-II motor activity. Instead, we find that spreading on soft substrates is impaired by reduced assembly of nascent, myosin-independent adhesions at the cell periphery. Enhancing integrin-ligand affinity through the addition of Mn^2+^ is sufficient both to stabilize nascent adhesions and increase cell spread area on soft substrates. We then implement a computational model to determine how changes in integrin-substrate catch bond kinetics affect integrin binding on substrates of different stiffness. We find that the biophysical properties of integrin-matrix catch bonds are optimized to sense changes in substrate stiffness around 6 kPa, consistent with our experimental results. Together these results illustrate that nascent adhesion formation in the lamellipodium functions as a myosin-II-independent mechanosensor to control cell adhesion and spreading.

## Results

### Spread area is a biphasic response of substrate stiffness, independent of myosin activity

To investigate the mechanisms that drive substrate stiffness sensing, we chose to measure the spread area of adherent cells. We first plated NIH 3T3 fibroblasts on a series of polyacrylamide gels covalently coupled with fibronectin, and with Young’s moduli ranging from 0.6 to 150 kPa (Fig. 1A). Cells were also plated on glass absorbed with fibronectin as a control. Consistent with previous reports (4, 6, 8-11), we find that cells’ spread area is sensitive to substrate stiffness (Fig. 1A). In contrast, however, we find that this response can be broken down into two regimes, there is poor spreading on soft ( < ∼5 kPa) substrates and high spreading on stiff ( > ∼8 kPa) substrates, with a transition region between these values and no statistical difference in spread area between populations within each regime (Fig. 1A). Furthermore, the morphology of cells on soft and stiff substrates is noticeably different (5). Cells on soft substrates are more rounded with disorganized actin cytoskeletons (Fig. 1B). In contrast, cells on stiff substrates exhibit more polarized shapes, and tend to have prominent stress fibers (Fig. 1B)

**Figure 1.**
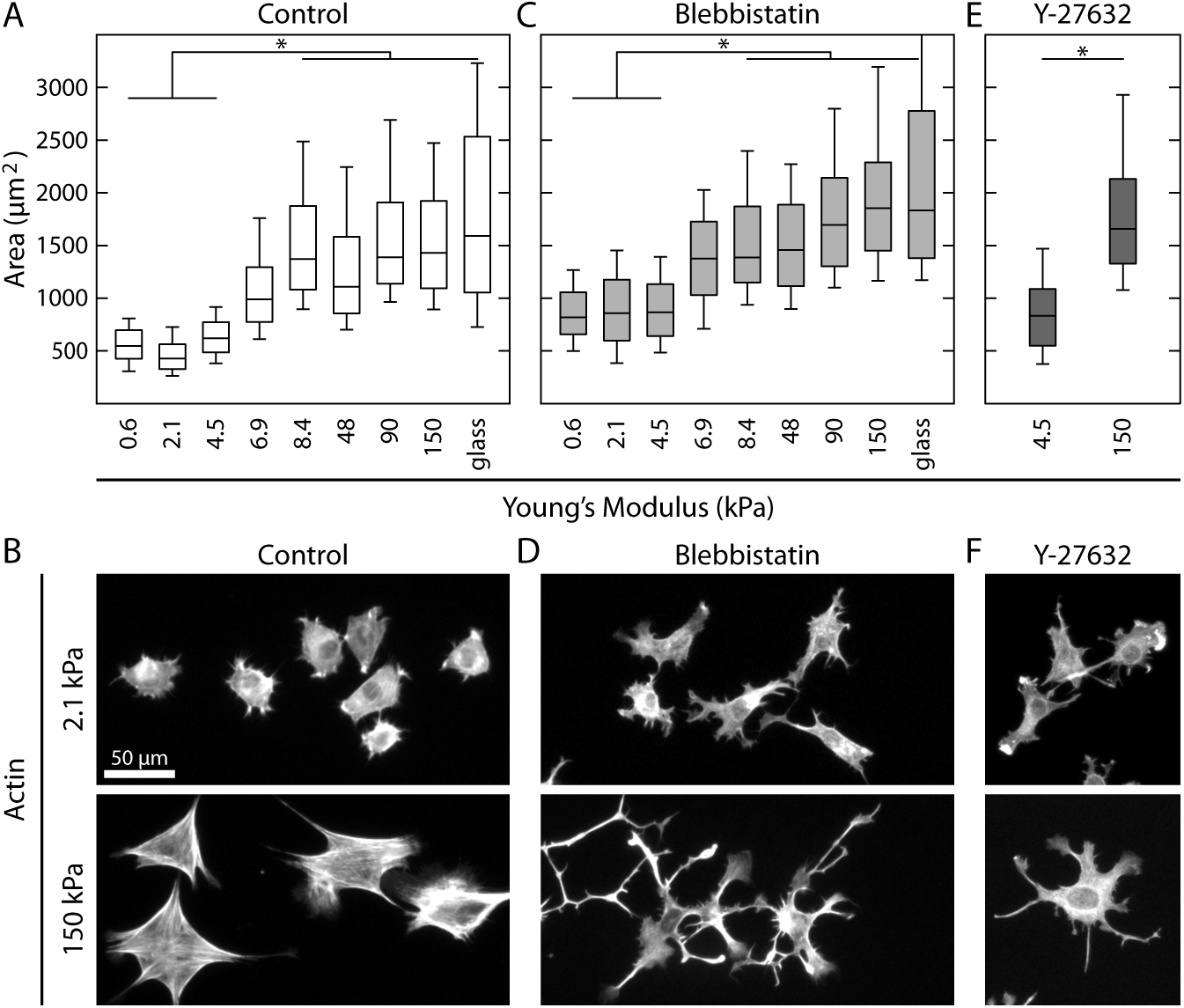
Spread area is a biphasic response of substrate stiffness, independent of myosin activity. (A) Boxplots of the spread area of NIH 3T3 fibroblasts plated on fibronectin coated polyacrylamide gels of varying stiffness. Cells can be grouped into soft (≤ 4.5 kPa) and stiff (≥ 8.4 kPa) regimes. From left to right N=182,674,205,155,254,400,205,487,170. (B) Representative images of control cells on soft and stiff substrates. (C) Boxplots of the spread area of cells treated with 50 μM blebbistatin to inhibit myosin activity. While blebbistatin treated cells spread more than control cells, they exhibit the same biphasic response as a function of substrate stiffness. From left to right N=169,228,329,67,159,125,119,183,56. (D) Representative images of blebbistatin treated cells on soft and stiff substrates. (E) Boxplots of the spread area of cells treated with with 20 μM Y-27632, which inhibits ROCK activity. Cells treated with Y-27632 still exhibit a difference in spread area on soft (N=148) and stiff (N=203) substrates. (F) Representative images of Y-27632 treated cells on soft and stiff substrates. Box plots represent 25^th^, 50^th^, and 75^th^ percentiles, while whiskers extend to the 10^th^ and 90^th^ percentiles. * indicates a p-value < 0.01.

Because myosin II activity has been widely implicated in mechanosensing (28), we next hypothesized that its inhibition would eliminate any change in spread area as a function of substrate stiffness. Surprisingly, cells incubated with 50 μM blebbistatin, a myosin II ATPase inhibitor, continue to exhibit a biphasic response to substrate stiffness (Fig. 1C). Cells treated with blebbistatin have an increased spread area compared to control cells across all stiffnesses, but exhibit the same soft and stiff regimes. Morphologically, myosin-inhibited cells on all substrates show more protrusions, but on stiff substrates the cells exhibit more spindle-like projections (Fig. 1D). Similar phenotypes are seen when cells are incubated with Rho-Kinase inhibitor (Y-27632; Fig. 1E,F), and when cells are plated on other ECM proteins (Fig. S1). Thus, the change in cell spread area that occurs between the soft and stiff regimes does not require myosin II activity.

### Substrate stiffness does not inhibit lamellipodia protrusion dynamics

To understand how substrate stiffness impacted cell spread area, we investigated the effects of substrate stiffness on protrusion dynamics. We tracked lamellipodia formation by taking time-lapse images of cells transiently transfected with a fluorescent membrane marker (GFP-stargazin) and treated with 20 μM Y-27632 on representative soft (2.1 kPA) and stiff (48 kPa) substrates 30 minutes after plating (Fig. 2A,B; Movies 1 and 2). Cells on soft substrates exhibit repeated cycles of protrusion and retraction, as seen in the kymograph (Fig. 2A), reducing their ability to spread. Cells on stiff substrates, however, exhibit continuous and steady protrusions that result in leading edge advance (Fig. 2B). Using cell contours derived from the fluorescence images we identified protrusive regions and measured their morphology and characteristics (Fig. 2C). We find no statistical difference between soft and stiff substrates for measurements of the average protrusion area (Fig. 2D) or the average protrusion width (Fig. 2E). These data indicate that substrate stiffness affects the stability of leading edge protrusions, but not the protrusion dynamics themselves. Arp2/3 mediated lamellipodium formation is still required for spreading, as cells on both soft and stiff substrates that are treated with CK-869, an Arp2/3 inhibitor, are indistinguishable from control cells on soft substrates (Fig. 2F,G). Together these results illustrate that it is the stabilization, not the formation, of Arp2/3-dependent lamellipodial protrusions that is hindered on soft substrates.

**Figure 2.**
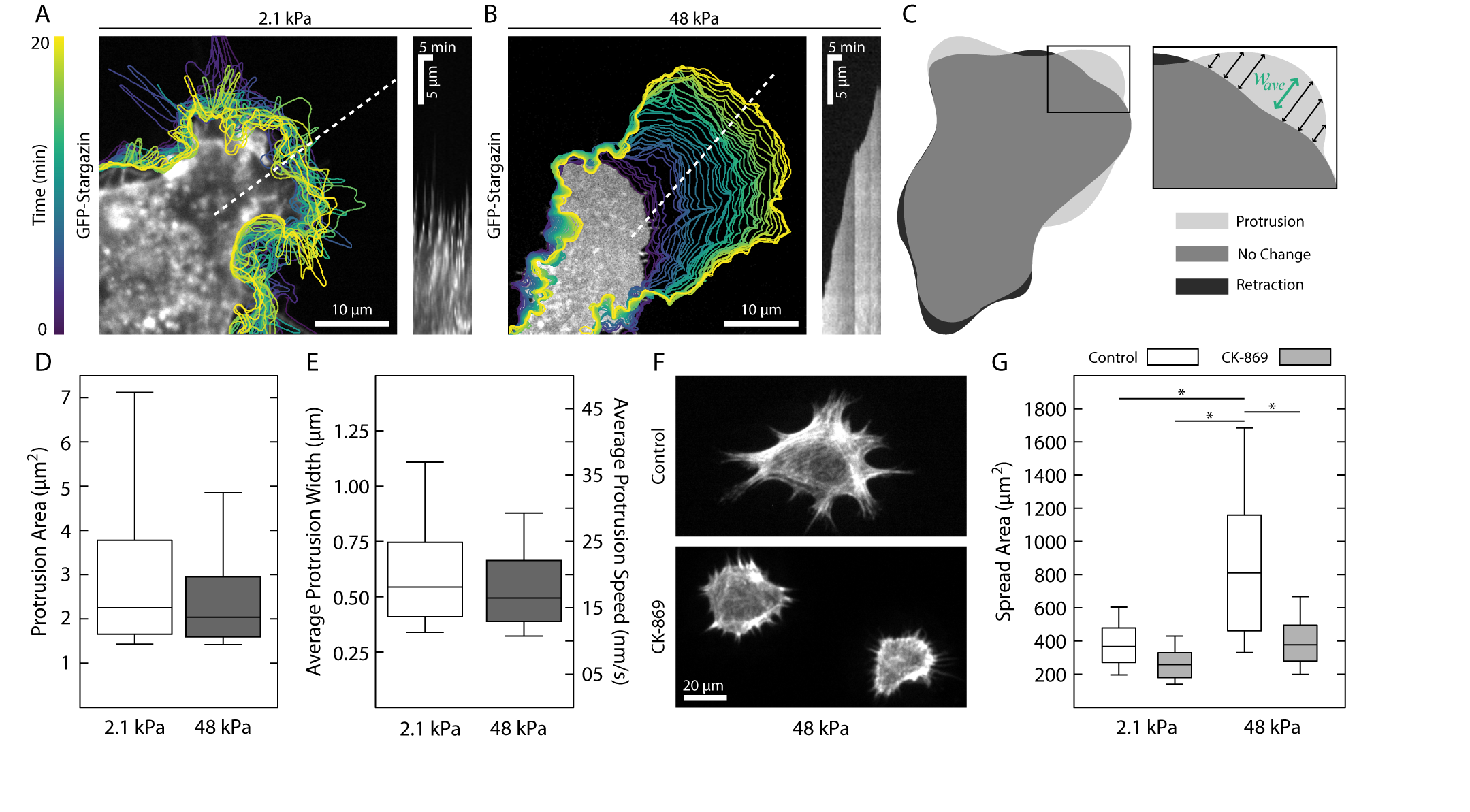
Soft substrates do not inhibit lamellipodia protrusion dynamics. (A-B) Contours of a cell expressing a GFP membrane marker plated on a soft and stiff substrates. On soft substrates protrusions are followed by rapid retractions, resulting in no advancement of the leading edge. In contrast, on stiff substrates, the leading edge advances continuously at each time step. The kymographs, taken along the dotted white lines, illustrate the different protrusion dynamics. (C) Protrusive regions were identified by overlaying successive contours and identifying new areas. The inset shows how the average width of the contour was calculated. (D) Boxplot showing the area of individual protrusions on soft (N=1722) and stiff (N=1800) substrates. No difference was seen between the two distributions. (E) The average protrusion width and speed were also indistinguishable between soft (N=1722) and stiff (N=1800) substrates. (F) Representative images of cells on stiff substrates treated with the arp2/3 inhibitor CK-869. Cells treated with CK-869 take on the morphology of control cells plated on soft substrates. (G) Boxplots of the spread area of cells treated with XX μM of CK-869. From left to right N=83,212,183,175. Box plots represent 25^th^, 50^th^, and 75^th^ percentiles, while whiskers extend to the 10^th^ and 90^th^ percentiles. * indicates a p-value < 0.01.

### Soft substrates impair nascent adhesion formation

To explore the mechanism of substrate stiffness-dependent changes in stabilization of myosin-II-independent protrusions, we examined the assembly of myosin-II-independent, nascent adhesions that form at the base of the lamellipodium. Two hours after plating, cells were treated with 20 μM Y-27632 for 30 min and then fixed and stained for actin, p34 (a subunit of Arp 2/3) and the focal adhesion protein paxillin (Fig. 3A,B). On both soft and stiff substrates p34 localizes to the cell periphery, indicative of the Arp2/3-dependent lamellipodium (Fig. 3A,B). On stiff substrates, paxillin forms small punctate nascent adhesions near the leading edge, which is characteristic of nascent adhesion formation on glass substrates (42). By contrast, on soft substrates, paxillin-rich nascent adhesions were seen at a lower density and formed further away from the leading edge. To quantify these differences in protein localization, we measured the average actin, p34 and paxillin intensity in ∼0.5 μm bands measured radially from the edge of the cell (Fig. 3C). We find that the peak of p34 intensity is localized right at the edge of the cell on all substrates. On stiff substrates, paxillin is located within ∼0.5 μm of the p34 peak (Fig. 3D). On soft substrates, there is a significantly reduced accumulation of paxillin, and its peak is found ∼5 μm behind the leading edge (Fig. 3D). These data suggest that cells have reduced nascent adhesion formation on soft substrates.

**Figure 3.**
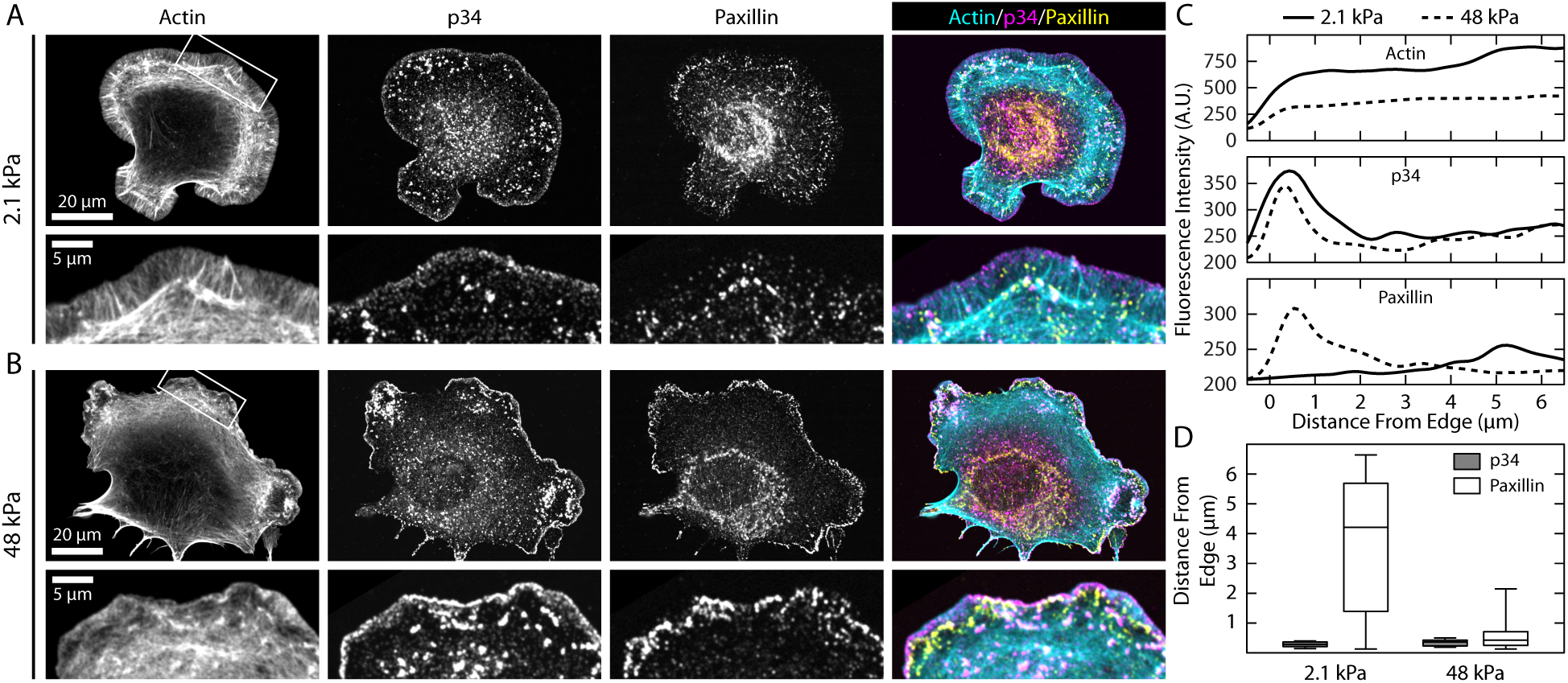
Soft substrates impair nascent adhesion formation. (A-B) Representative immunofluorescence images of cells on soft and stiff substrates showing actin, p34 (a subunit of arp2/3), and paxillin (focal adhesion protein). Cells on soft substrates exhibit large lamellipodia and reduced paxillin staining. No noticeable difference can be seen in the p34 localization. (C) Average linescans for each of the three channels for the regions shown in the insets of A-B. While p34 shows a peak at the same position on both soft and stiff substrates, the peak for paxillin is much further behind the leading edge, and much less intense. (D) Boxplots showing the distance from the leading edge for both the p34 (grey) and paxillin (white) signal on soft (N=46) and stiff (N=42) substrates. Box plots represent 25^th^, 50^th^, and 75^th^ percentiles, while whiskers extend to the 10^th^ and 90^th^ percentiles.

### Activation of integrins via Mn^2+^ is sufficient to promote spreading on soft substrates

Given the reduced density of nascent adhesions on soft substrates, we sought to explore the extent to which changes in integrin-ligand affinity could stimulate their formation. The presence of 3 μM Mn^2+^ increases the lifetime of integrin-fibronectin bonds (43). When cells were plated on soft substrates in the presence of 3 μM Mn^2+^ they exhibited a greater than two-fold increase in spread area on soft substrates, similar to their spread area on stiff substrates either in the presence and absence of Mn2+ (Fig. 4A). To directly compare the effect of Mn^2+^ on adhesion assembly on soft substrates, we performed immunofluorescence of paxillin and actin. Addition of Mn^2+^ to cells on soft substrates stimulated the formation of paxillin-rich adhesions near the cell periphery and even the formation of lamellar actin bundles.

**Figure 4.**
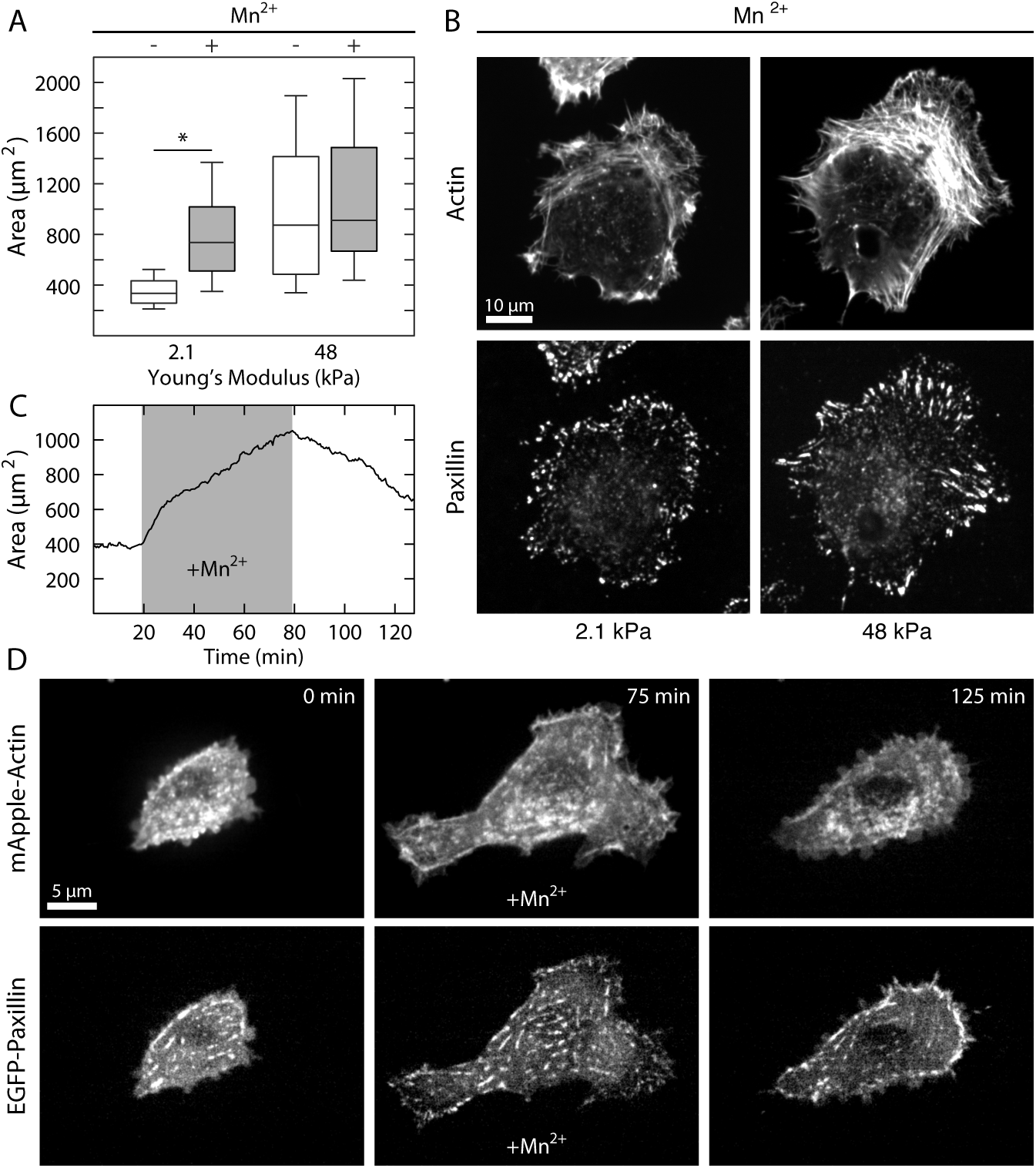
Activation of integrins via Mn^2+^ is sufficient to promote spreading on soft substrates. (A) Boxplots of spread area for control cells and cells treated with 3 μM of Mn^2+^ on soft and stiff substrates. Cells treated with Mn^2+^ on soft substrates were not significantly different from control cells on stiff substrates. From left to right N=512,634,300,216. (B) Representative immunofluorescence images of cells treated with Mn^2+^ on soft substrates and control cells on stiff substrates. The cells treated with Mn^2+^ on soft substrates take on the morphology of control cells on stiff substrates. (C) Plot of area vs time for a cell on soft substrate. 3 μM Mn^2+^ was flowed into the imaging chamber at ∼20 min, and then washed out again at ∼80 min. As soon as the Mn^2+^ is added to the solution, the cell begins to spread. When the Mn^2+^ is washed out, the cell begins to retract. (D) Representative images from the Mn^2+^ wash-in time course shown in C. After addition of Mn^2+^ both focal adhesions and actin stress fibers can be seen to form. Box plots represent 25^th^, 50^th^, and 75^th^ percentiles, while whiskers extend to the 10^th^ and 90^th^ percentiles. * indicates a p-value < 0.01.

To determine how rapidly Mn^2+^ could induce changes in adhesion formation and cell spread area, we performed live cell imaging of EGFP-paxillin and mApple-Actin in cells plated on a soft substrate during addition of Mn^2+^ to the media. (Fig. 4C,D; Movie 3). Prior to addition of Mn2+, there is significant protrusive activity on soft substrates, but no change in area or cell shape. Upon addition of 3 μM Mn^2+^, protrusions stabilized, new focal adhesions formed and the cell increased in spread area (Fig. 4C,D; Movie 3). After an hour of incubation the media was again replaced with media lacking Mn^2+^, and the cell immediately began to retract back towards its initial spread area (Fig. 4C,D; Movie 3). Thus the presence of Mn^2+^ is sufficient to promote spreading on soft substrates. This strongly suggests that integrin-fibronectin bond affinity plays an important role in substrate stiffness sensing to mediate cell spreading.

### Integrin catch-bond kinetics mediate substrate stiffness sensing

To explore how integrin-fibronectin binding kinetics could enable substrate stiffness sensing we built a computational model of nascent adhesion assembly at the leading cell edge. The model, similar to previous approaches (23-27), incorporates biophysical properties of cell-matrix adhesions, actin retrograde flow and substrate rigidity. Individual integrins in the model act as molecular clutches, intermittently transmitting force produced by actin retrograde flow to the substrate. It has been shown previously, that integrin-fibronectin bonds are catch-bonds, meaning their lifetime increases as a function of load (43). We used our model to explore which features of these bond kinetics are important in mediating substrate stiffness sensing in nascent adhesions.

In the model, both integrins and ligands are represented as single point particles. Initially, a given number of fibronectin molecules are randomly attached to a substrate, to which integrins can bind. The integrins undergo cycles of diffusion, binding, and unbinding along a quasi-2D surface mimicking the ventral membrane of cells, above the substrate (Fig. 5A,B). Integrins bound to actin undergo retrograde flow, as is seen in the lamellipodium (44), while unbound integrins are free to diffuse on the surface (45). When an integrin comes in close proximity of a free fibronectin, it establishes a harmonic potential interaction, which mimics binding, with the stiffness determined by substrate rigidity. The assumption of simultaneous binding of the integrin to both the substrate ligand and the actin is motivated by the need to build tension on the integrin-fibronectin bond and this tension regulates the bond lifetime. By keeping a constant actin flow, forces on the bonds are directly proportional to the substrate stiffness. All parameters in the model are based upon available experimental data (37, 45-47) (See Methods). In particular, we directly incorporated the lifetime versus force relationships of integrin-fibronectin bonds from AFM single molecule experiments (43). To quantify the amount of integrin binding, we measured the average fraction of bound integrins over the course of the simulations (between 10-300 s) for each condition.

**Figure 5.**
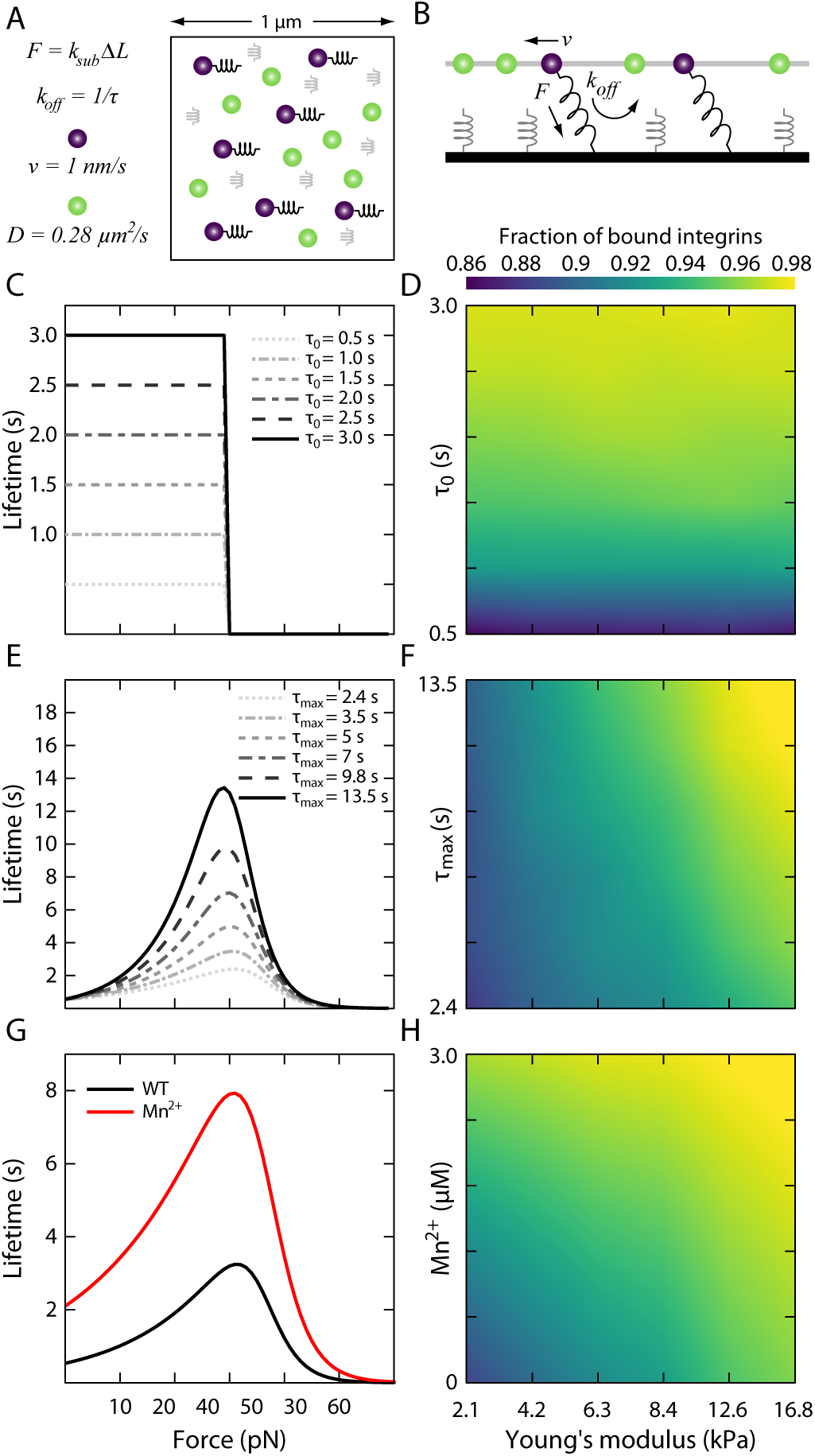
Computational model of integrin-based adhesion dynamics. (A-B) Schematic of the computational model from the top (A) and side (B) perspectives. Two quasi-2D surfaces are placed 200 nm apart. The bottom surface represents the substrate and consists of a bundle of ideal springs with stiffness, *k*_*sub*_, proportional to the substrate Young’s modulus. The top surface mimics a representative unit of the central surface of a fibroblast, with integrins diffusing (green particles) with diffusion coefficient, *D*, and establishing interactions with the substrate springs (purple particles). Upon binding the substrate, a force is exerted on the integrin particle parallel to the substrate and builds tension on the bond, which determines the integrin unbinding rate, *k*_*оƒƒ*_. (C) Relations of lifetime versus tension with unloaded lifetimes equal to the maximum lifetime, *τ*_*0*_ = *τ*_*max*_ at tensions **<** 30 pN and zero otherwise. (D) The corresponding average fraction of ligand bound integrins as a function of varying the substrate’s Young’s modulus. The average fraction of bound integrins is insensitive to substrate stiffness. (E) Relations of lifetime versus tension with fixed *τ*_*0*_ and increasing *τ*_*max*_. (F) The corresponding average fraction of bound integrins in this case is only weakly sensitive to substrate stiffness. (G) Lifetime versus tension relationship for WT and Mn^2+^ treated integrins, which amounts to both a shift in *τ*_*0*_ and in *τ*_*max*_. (H) The corresponding average fraction of bound integrins. The number of bound integrins on soft substrates in the presence of Mn^2+^ is identical to the number of bound integrins for WT integrins on stiff substrates.

We first tested whether catch-bond behavior is required for substrate stiffness sensing by simulating the lifetime versus force relationship as a step-function. The bond lifetime was held constant below a peak force of 30 pN, and zero for higher forces (Fig. 5C). Varying the magnitude of the integrin-fibronectin lifetime has no effect on the fraction of bound integrins as a function of substrate stiffness (Fig. 5D). Therefore, in the absence of a force-dependent catch-bond mechanism, integrin binding kinetics are independent of substrate rigidity. We next explored how changes in catch-bond biophysical properties engendered stiffness sensing. We simulated the lifetime versus force relationship as a typical catch-bond, and varied the maximum lifetime for the 30 pN peak force, keeping the unloaded lifetime constant (Fig. 5E). In these conditions, because the integrin-fibronectin bond lifetime increases with increasing force, and force is modulated by substrate stiffness, actin flow enhances the amount of bound integrins on stiff substrates (Fig. 5F). Changes in the maximum lifetime of the bond, however, have little impact on the fraction of bound integrins for a given substrate stiffness (Fig. 5F). Surprisingly, the transition between different regimes of integrin binding using known biophysical parameters of integrin catch bonds occurs in the model naturally around a Young’s modulus of ∼7 kPa, as is seen in our experiments (Fig. 1).

In the presence of Mn^2+^, integrin-fibronectin bonds have both an increased affinity and enhanced bond lifetime in response to increased tension with respect to wild type conditions (43, 48-50). To mimic these effects in the model, we combined our two previous results, increasing both the unloaded lifetime and the lifetime at the peak force (Fig. 5G). By increasing the integrin-fibronectin bond lifetime for forces lower than 30 pN, actin flow enhances the amount of bound integrins on soft substrates (Figure 5H). Under these conditions the fraction of bound integrins on soft substrates in the presence of Mn^2+^ is quantitatively similar to the fraction of bound integrins on stiff substrates in control conditions. Thus, while neither an increase in unloaded lifetime or lifetime at peak force is sufficient on its own to recapitulate the effects of Mn^2+^, their combined effect is enough to abrogate the effects of substrate stiffness on cell spreading.

Collectively these results illustrate that rigidity sensing in the lamellipodium is determined by catch-bond kinetics of integrin-fibronectin bonds, and that the fraction of bound integrins is sensitive to both the unloaded lifetime and the maximum lifetime of the catch-bond curve. Addition of Mn^2+^ results in longer integrin lifetimes on soft substrates, thereby increasing the average fraction of bound integrins. This change in integrin binding kinetics allows cells to spread on soft substrates.

## Discussion

The ability of cells to sense the stiffness of their extracellular environment is critical to their ability to regulate growth, viability, migration and differentiation (1-3). Here we demonstrate that fibroblasts exhibit a biphasic response in spreading on matrices of variable stiffness (Fig. 1). For matrices with a Young’s modulus less than ∼5 kPa, cells are poorly spread with minimal adhesion assembly and few organized actin structures. Above ∼8 kPa, fibroblasts achieve a maximal spread area with typical adhesion assembly and highly organized actin cytoskeletons. This transition stiffness is comparable to physiological tissue stiffness (16), and is of the same order of magnitude as values reported previously (8-12). While it has been suggested that cell spread area as a function of substrate stiffness follows a power-law behavior (9, 12), using a larger number of substrates we find that it is better described as biphasic.

Due to its overwhelming role in cellular force generation, myosin II has been presumed to be the predominant mechanism of substrate stiffness sensing by adherent cells (23-28). Here, however, we demonstrate a myosin-independent stiffness sensing mechanism that controls spread area and arises from forces generated by actin polymerization within the lamellipodium. Integrins, which connect and transmit stress between the cytoskeleton and the extracellular matrix, behave as catch-bonds whose lifetime is determined as a function of the applied load (43, 51). As the load on the integrin increases, the lifetime of the bond also increases (43). On stiff substrates, this increase in lifetime is sufficient to promote clustering and adhesion formation (Fig. 4). Conversely, on soft substrates the reduction in stiffness leads to shorter bond lifetimes which inhibit the required clustering for adhesion formation (Fig. 5). Both our experimental and simulation data suggest that integrin force-dependent binding kinetics are most sensitive to substrates with a stiffness between approximately 5-8 kPa. Addition of Mn^2+^, which alters the kinetics of integrin-ECM bonds by increasing the unloaded and peak force lifetimes (43, 48-50), both increases the number of bound integrins, and decreases the average spacing between bound integrins. Together these effects promote adhesion formation and enable cells on soft substrates to spread and take on the morphology characteristics of cells on stiff substrates (Fig. 4).

Conceptually, this framework is similar to the general motor-clutch model (52) that has been previously suggested as a mechanism for understanding mechanosensitivity (23, 24, 26, 27). Instead of forces being generated by myosin motors, the force applied across the integrin bonds is generated by actin polymerization in the lamellipodium. These polymerization forces modulate the integrin-ECM bond kinetics and offer a surprisingly simple and elegant mechanism to understand substrate stiffness sensing. Previous work has established that there is a minimum spacing required between integrins for adhesion formation (38). Binding of integrins to their ligands also limits their diffusion in the membrane (45) and drives clustering at the nanoscale (36). Once a nanoscale cluster of integrins has formed, the force required to rupture the adhesion, i.e. the adhesion strength, is more than an order of magnitude greater than typical tensions generated in the cytoskeleton (53). Thus by increasing the density of bound integrins, adhesion stabilization is increased and the cell is able to spread.

Together these results suggest that the lamellipodium acts as a myosin independent mechanosensor, applying force to bound integrins via actin polymerization driven retrograde flow. On soft substrates, the increased pliability of the matrix leads to a reduced load on the integrin-ECM bond, resulting in a shorter lifetime. This shorter lifetime, prevents integrin clustering and thereby inhibits adhesion stabilization, leading to a poor ability to spread. On stiff substrates, integrin-ECM bonds experience greater loads and thus increased lifetimes, which promote adhesion stabilization and enable cells to spread. While these results do not exclude the possibility that myosin generated forces may be one mechanism to probe substrate stiffness, they suggest that stiffness sensing emerges passively from the properties of the integrin binding kinetics. Given that simply shifting these kinetics can induce spreading on soft substrates, it will be interesting in the future to explore whether this approach is sufficient to recover other functions found to be impaired by soft substrates, as in development, differentiation and disease

## Acknowledgements

This research was supported through internal funds from the University of Rochester to P.W.O, NIGMS GM085087 to M.L.G. and the DOD/ARO through a MURI grant W911NF1410403 to M.L.G and G.A.V. This work was also partially supported by the University of Chicago Materials Research Science and Engineering Center, which is funded by the National Science Foundation under award number DMR-1420709. The National Science Foundation XSEDE resources at the Pittsburgh Supercomputing Center provided computational time.

## Author Contributions

P.W.O and M.L.G designed the research. P.W.O., Y.B, G.R.R.-S.J. and S.P.W. performed the experiments. T.C.B. and G.A.V. designed the model. T.C.B. performed the simulations. P.W.O., T.C.B. and A.V.S. analyzed the data. P.W.O, T.C.B. and M.L.G. wrote the paper.

## Materials and Methods

### Cell culture and reagents

NIH 3T3 mouse fibroblasts (American Type Culture Collection) were cultured in DMEM media (Mediatech, Inc.) supplemented with 10% FBS (HyClone; Thermo Fisher Scientific), 2 mM l-glutamine (Invitrogen), and penicillin-streptomycin (Invitrogen). Cells were tested for mycoplasma and were free of contamination. Cells were transiently transfected with a plasmid DNA construct encoding GFP-Stargazin (a gift from A. Karginov, University of Illinois at Chicago, Chicago, IL). The following antibodies were used: mouse anti-Pxn and rabbit anti-p34 (Millipore); Cy5 donkey anti– mouse (Jackson ImmunoResearch Laboratories, Inc.); Alexa Fluor 568 goat anti–rabbit (Invitrogen). Alexa Fluor 488 phalloidin was purchased from ThermoFisher Scientific. Blebbistatin was purchased from Sigma and used at 50 μM. The Rho Kinase inhibitor Y-27632 was purchased from EMD Millipore and used at 20 μM. The ARP2/3 inhibitor CK-869 and control compound CK-312 were purchased from Calbiochem and used at 50 μM. The RGDs cyclo (Arg-Gly-Asp-D-Phe-Lys) and H-Arg-Gly-Asp-Ser-Lys-OH1 were purchased from Peptides International. Fibronectin derived from human plasma was purchased from Millipore. Vitronectin and Collagen were purchased from Thermo Fisher Scientific. Mn^2+^ was purchased from Fischer Scientific and used at 3 μM.

### Polyacrylamide substrates

PAA substrates were prepared on glass coverslips using previously published methods (54, 55). In brief, acrylamide/ bis-acrylamide were used to create PAA gels with Young’s moduli of 0.6, 2.1, 4.5, 6.9, 8.4, 48, 90, and 150 kPa (5, 55). Fibronectin, collagen, and RGDs were coupled to the surface of the PAA gels using the photoactivatable crosslinker sulfo-SANPAH (Thermo Fisher Scientific). PAA gels were covered with a 2.5 mg/ml solution of sulfo-SANPAH and exposed to an 8-W UV lamp for 5 min. The PAA gels were rinsed with PBS and incubated with 1 mg/ml fibronectin or RGD at room temperature for 45 min, or in 2 mg/ml collagen at 4˚C overnight. The PAA gels were then rinsed repeatedly and plated with cells. Vitronectin (ThermoFisher Scientific) was coupled to the surface of the PAA gels EDC/NHS chemistry (32, 56). Briefly, the polymerized gel was placed in a UVO-Cleaner 342 (Jelight, Irvine, CA) and illuminated with 185- and 254-nm ultraviolet light for 90 s. Gels were incubated in 200 μl of a solution containing 5 mg/mL EDC (Thermo Fisher Scientific) and 10 mg/mL NHS (Thermo Fisher Scientific) for 20 min. The EDC-NHS solution was then aspirated and replaced with a solution containing 10 μg/mL vitronectin in a buffer of HEPES (pH 8.5) for 20 min. Gels were washed 3 times for 5 min in phosphate-buffered saline (PBS) before cells were plated. Cells were plated on substrates for 2.5 hours. CK compounds were added at plating and Blebbistatin, Y-27632 and Manganese were added after 2 hours unless otherwise noted.

### Microscopy and live cell imaging

Images were obtained using Metamorph (Molecular Devices) acquisition software on either an Nikon Ti-E inverted microscope equipped with a metal halide light source (Lumen 200PRO; Prior Scientific) or a Nikon Eclipse Ti-E inverted microscope with a Yokogawa CSU-X1 confocal scanhead and Spectral Applied Research Laser Merge Module (491 nm, 561 nm, and 643 nm Lasers). Images were obtained with a Photometrics Coolsnap HQ2 CCD camera using 20x Plan Fluor ELWD 0.45 NA, 40x Plan Fluor 1.3 NA, or 60X Plan Apo 1.2 NA objectives (Nikon). For live cell imaging, cells were mounted in a perfusion chamber (Warner Instruments) and maintained at 37˚C. Media for live cell imaging was supplemented with 10 mM HEPES and 30 mL/mL Oxyrase (Oxyrase Inc.).

### Immunofluorescence

Cells were rinsed in warm cytoskeleton buffer (10 mM MES, 3 mM MgCl2, 1.38 M KCl, and 20 mM EGTA) and then fixed and permeabilized in 4% PFA (Electron Microscopy Sciences), 1.5% bovine serum albumin (Thermo Fisher Scientific), and 0.5% Triton X-100 in cytoskeleton buffer for 15 min at 37C. Gels were then rinsed three times in PBS and incubated with mouse anti-paxillin and rabbit anti-p34 (1:400; Millipore) for 1 h at room temperature. The gels were then rinsed three times in PBS and before being incubated with fluorescently labeled secondary antibodies and phalloidin. Gels were rinsed three times and mounted in non-curing media (SlowFade; Invitrogen) and sealed with nail polish.

### Image Analysis

All image analysis was done using ImageJ, MATLAB (Mathworks) or Python. Cell area was determined by thresholding images of actin to create binary masks. Protrusion analysis was performed by thresholding each image in a time series to create a binary mask, from which the cell contour could be extracted. Protrusive regions were identified by overlaying successive contours to identify regions of new area. Each protrusion was segmented to identify the total area, along with the leading and trailing edges. A given protrusion width was determined by taking each point along the leading edge contour and determining the nearest distance to the trailing edge contour, and averaging across the entire set. The protrusion speed was calculated by dividing the average width by the frame interval of the time series.

To determine the location of intensity maxima for p34 and paxillin, images were first thresholded in the actin channel as described above to create a cell contour. We then performed a succession of 1 pixel erosions, to create a series of contours that radially propagated towards the center of the cell. Lamellapodia regions were then identified and masked. To create linescans, we averaged the intensity signal in a window that was 5 contours in width (∼ 0.5 μm) and shifted the window 1 contour towards the center of the cell for each step. This results in an average radial intensity from the edge of the cell. The distance plotted is the distance from the edge of the cell to the center of the window.

## Computational Model

We use a Brownian dynamics approach to simulate integrin binding and unbinding in response to differences in the substrate stiffness. Integrins are represented as single point particles that undergo cycles of diffusion, binding, and unbinding, along a quasi-2D surface mimicking the ventral surface of cells. The substrate is represented as an isotropic and elastic material, consisting of a bundle of ideal springs which mimic ligands. In order to simulate the effect of Mn^2+^ on integrin binding, we modulate the relationship of bond lifetime versus force.

### Computational domain and boundary conditions

The computational domain is 3D and consists of two parallel surfaces: the bottom layer is fixed in space and represents the substrate (Fig 5B); the top layer is an ideal surface, where integrin particles diffuse along x and y, with a diffusion coefficient of *D* = 0.28 *μm*^*2*^*/s* (45). Integrin particles are harmonically restrained in the vertical direction, with an equilibrium distance, *L*, of 20 nm from the bottom layer. This distance between the integrin layer and substrate corresponds to the separation of the open extracellular integrin headpiece from the membrane (46). From the top, the computational domain is a square with side dimensions of 1 μm (Figure 5A). In order to avoid finite size effects on integrin motion, we use periodic boundary conditions in the lateral directions, x and y.

### Substrate model

The substrate is an elastic solid, consisting of a bundle of ideal linear springs, with stiffness depending on the substrate rigidity, as:

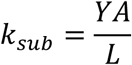

where *Y* is the Young’s modulus (we tested values in the range 2.1-16.8 kPa), *A* is the integrin/ligand cross-sectional area (corresponding to 80 nm, from an ideal bar of radius ∼5nm, corresponding to approximately half the value of an integrin transmembrane legs separation), and *L* is the equilibrium distance separation between substrate and top layer.

Hooke’s law for each spring in the bundle can be written as:

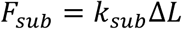

with *ΔL* corresponding to the deviation from the equilibrium distance separation between the two surfaces. We use a spring density of 200 per μm^2^, however our results, which are expressed in terms of average fraction of bound integrins at the steady state, do not depend on the density of substrate springs.

### Integrin particles and implementation algorithm

Integrin particles on the top surface diffuse in Brownian motion. At each time step of the simulation, the positions of each integrin particle, /’, is updated, according to the Langevin equation of motion, with inertia negligible:

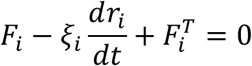

We use a time step *dt* = 10−^4^ *s* and a friction ζ = 0.0142 *pN s/μm*, corresponding to a diffusion coefficient *D* = 0.28 *μm*^*2*^*/s* (45), from:

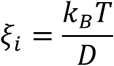

with *k*_*B*_*T* = 4.11 *pN* · *nm*.

The force acting on integrin particles in the Langevin equation of motion has two contributions: a deterministic contribution and a stochastic contribution. The deterministic contribution comes from the tension of the bond towards the substrate and from the imposed velocity along *xy*:

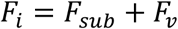

The stochastic contribution represents thermal fluctuations and obeys the fluctuation-dissipation theorem (57):

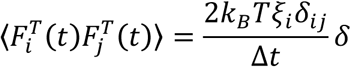

where *δ*_*ij*_ is the Kronecker delta, and *δ* is a second-order tensor.

To integrate over time and update the positions of the various elements in the simulation, we use the explicit Euler integration scheme:

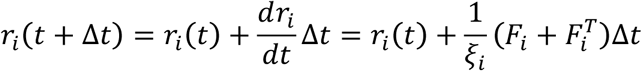

Integrins can establish harmonic interactions with substrate springs, when in proximity of them, if the spring is free from a previous bond. Upon binding the substrate, integrin particles are subjected to a force parallel to the substrate, *F_v_*, corresponding to a velocity of 1 nm/s, of the order of lamellipodium actin polymerization (44, 58).

The integrin/substrate interaction persists for a characteristic lifetime, τ, which depends upon the tension on the bond and follows the formalism of the implemented integrin unbinding rate, *k*_*оƒƒ*_.

A double exponential pathway determines unbinding rates, as a function of tension, *ƒ*. It includes a strengthening pathway, with a negative exponent, and a weakening pathway, with a positive exponent. For wild type conditions, the unbinding rate is:

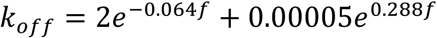

For Mn^2+^ coditions, the unbinding rate is:

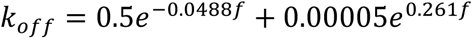

The functional form of the catch bonds is taken from a model that assumes a single bound state and two unbinding pathways (59, 60) and was previously used for integrin-based adhesions (61). The coefficients of this form are estimated for reproducing maximum bond lifetime and corresponding tension from (43) and integrin unloaded affinity (48-50).

---

**Supplementary Figure S1.**
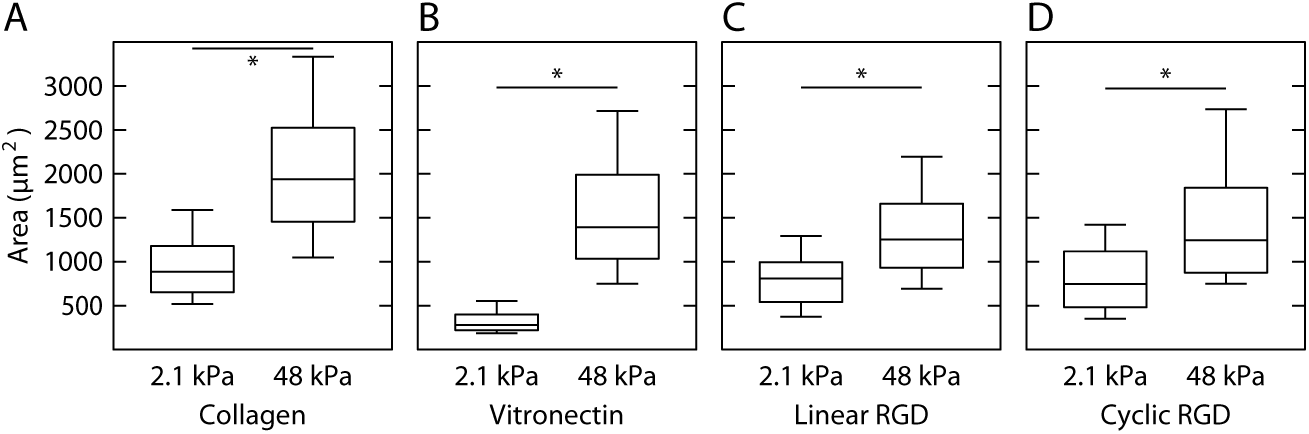
Cells spread area in response to substrate stiffness is independent of ECM ligand. (A) Cells were plated on soft (N=240) and stiff (N=255) substrates coated with collagen. (B) Cells were plated on soft (N=74) and stiff (N=55) substrates coated with vitronectin. (C) Cells were plated on soft (N=137) and stiff (N=150) substrates coated with linear RGD peptide. (D) Cells were plated on soft (N=145) and stiff (N=69) substrates coated with cyclic RGD peptide.

## Movie 1

An NIH 3T3 fibroblast expressing a GFP membrane marker plated on a soft (2.1 kPa Young’s modulus) substrate in the presence of 20 μM Y-27632. Time is in min:sec. From Fig. 2A.

## Movie 2

An NIH 3T3 fibroblast expressing a GFP membrane marker plated on a stiff (48 kPa Young’s modulus) substrate in the presence of 20 μM Y-27632. Time is in min:sec. From Fig. 2B.

## Movie 3

An NIH 3T3 fibroblast expressing mApple-actin and EGFP-paxillin plated on a soft (2.1 kPa young’s modulus) substrate. AT ∼20 min 3μM Mn^2+^ is flowed into the imaging chamber. At ∼80 min the Mn^2+^ is washed out of the chamber. Time is in hr:min:sec. From Fig. 4C-D.

## References

1. Paluch EK, et al. (2015) Mechanotransduction: use the force(s). BMC Biol 13(1):47.

2. Iskratsch T, Wolfenson H, Sheetz MP (2014) Appreciating force and shape—the rise of mechanotransduction in cell biology. Nat Rev Mol Cell Biol 15(12):825–833.

3. Charras G, Sahai E (2014) Physical influences of the extracellular environment on cell migration. Nat Rev Mol Cell Biol 15(12):813–824.

4. Lo C-M, Wang H-B, Dembo M, Wang Y-L (2000) Cell movement is guided by the rigidity of the substrate. Biophys J 79(1):144–152.

5. Yeung T, et al. (2005) Effects of substrate stiffness on cell morphology, cytoskeletal structure, and adhesion. Cell Motil Cytoskeleton 60(1):24–34.

6. Prager-Khoutorsky M, et al. (2011) Fibroblast polarization is a matrix-rigidity-dependent process controlled by focal adhesion mechanosensing. Nat Cell Biol. doi:10.1038/ncb2370.

7. Oakes PW, et al. (2009) Neutrophil morphology and migration are affected by substrate elasticity. Blood 114(7):1387–1395.

8. Califano JP, Reinhart-King CA (2010) Substrate Stiffness and Cell Area Predict Cellular Traction Stresses in Single Cells and Cells in Contact. Cell Mol Bioeng 3(1):68–75.

9. Han SJ, Bielawski KS, Ting LH, Rodriguez ML, Sniadecki NJ (2012) Decoupling Substrate Stiffness, Spread Area, and Micropost Density: A Close Spatial Relationship between Traction Forces and Focal Adhesions. Biophys J 103(4):640–648.

10. Ghibaudo M, et al. (2008) Traction forces and rigidity sensing regulate cell functions. Soft Matter 4(9):1836–1843.

11. Tee S-Y, Fu J, Chen CS, Janmey PA (2011) Cell shape and substrate rigidity both regulate cell stiffness. Biophys J 100(5):L25–7.

12. Engler A, et al. (2004) Substrate compliance versus ligand density in cell on gel responses. Biophys J 86(1 Pt 1):617–628.

13. Breckenridge MT, Desai RA, Yang MT, Fu J, Chen CS (2014) Substrates with engineered step changes in rigidity induce traction force polarity and durotaxis. Cell Mol Bioeng 7(1):26–34.

14. Raab M, et al. (2012) Crawling from soft to stiff matrix polarizes the cytoskeleton and phosphoregulates myosin-II heavy chain. J Cell Biol. doi:10.1083/jcb.201205056.

15. Wang HB, Dembo M, Wang YL (2000) Substrate flexibility regulates growth and apoptosis of normal but not transformed cells. Am J Physiol Cell Physiol 279(5):C1345–50.

16. Engler A, Sen S, Sweeney H, Discher D (2006) Matrix elasticity directs stem cell lineage specification. Cell 126(4):677–689.

17. Fu J, et al. (2010) Mechanical regulation of cell function with geometrically modulated elastomeric substrates. Nat Methods 7(9):733–736.

18. Paszek MJ, et al. (2005) Tensional homeostasis and the malignant phenotype. Cancer Cell 8(3):241–254.

19. Mekhdjian AH, et al. (2017) Integrin-mediated traction force enhances paxillin molecular associations and adhesion dynamics that increase the invasiveness of tumor cells into a three-dimensional extracellular matrix. Mol Biol Cell:mbc.E16–09–0654.

20. Zaidel-Bar R, Itzkovitz S, Ma’ayan A, Iyengar R, Geiger B (2007) Functional atlas of the integrin adhesome. Nat Cell Biol 9(8):858–867.

21. Schiller HB, Friedel CC, Boulegue C, Fässler R (2011) Quantitative proteomics of the integrin adhesome show a myosin II-dependent recruitment of LIM domain proteins. EMBO Rep 12(3):259–266.

22. Kanchanawong P, et al. (2010) Nanoscale architecture of integrin-based cell adhesions. Nature 468(7323):580–584.

23. Chan CE, Odde DJ (2008) Traction dynamics of filopodia on compliant substrates. Science 322(5908):1687–1691.

24. Macdonald A, Horwitz A, Lauffenburger D (2008) Kinetic model for lamellipodal actin-integrin “clutch” dynamics. Cell Adh Migr 2(2):95–105.

25. Bangasser BL, Odde DJ (2013) Master equation-based analysis of a motor-clutch model for cell traction force. Cell Mol Bioeng 6(4):449–459.

26. Elosegui-Artola A, et al. (2014) Rigidity sensing and adaptation through regulation of integrin types. Nat Mater 13(6):631–637.

27. Elosegui-Artola A, et al. (2016) Mechanical regulation of a molecular clutch defines force transmission and transduction in response to matrix rigidity. Nat Cell Biol 18(5):540–548.

28. Plotnikov SV, Pasapera AM, Sabass B, Waterman CM (2012) Force Fluctuations within Focal Adhesions Mediate ECM-Rigidity Sensing to Guide Directed Cell Migration. Cell 151(7):1513–1527.

29. Yan J, Yao M, Goult BT, Sheetz MP (2015) Talin Dependent Mechanosensitivity of Cell Focal Adhesions. Cell Mol Bioeng 8(1):151–159.

30. Thievessen I, et al. (2013) Vinculin-actin interaction couples actin retrograde flow to focal adhesions, but is dispensable for focal adhesion growth. J Cell Biol 202(1):163–177.

31. Sawada Y, et al. (2006) Force sensing by mechanical extension of the Src family kinase substrate p130Cas. Cell 127(5):1015–1026.

32. Oakes PW, Banerjee S, Marchetti MC, Gardel ML (2014) Geometry regulates traction stresses in adherent cells. Biophys J 107(4):825–833.

33. Hu K, Ji L, Applegate KT, Danuser G, Waterman-Storer CM (2007) Differential transmission of actin motion within focal adhesions. Science 315(5808):111–115.

34. Aratyn-Schaus Y, Gardel ML (2010) Transient frictional slip between integrin and the ECM in focal adhesions under myosin II tension. Curr Biol 20(13):1145–1153.

35. Gardel ML, et al. (2008) Traction stress in focal adhesions correlates biphasically with actin retrograde flow speed. J Cell Biol 183(6):999–1005.

36. Bachir AI, et al. (2014) Integrin-associated complexes form hierarchically with variable stoichiometry in nascent adhesions. Curr Biol 24(16):1845–1853.

37. Alexandrova AY, et al. (2008) Comparative dynamics of retrograde actin flow and focal adhesions: formation of nascent adhesions triggers transition from fast to slow flow. PLoS ONE 3(9):e3234.

38. Cavalcanti-Adam E, et al. (2007) Cell spreading and focal adhesion dynamics are regulated by spacing of integrin ligands. Biophys J 92(8):2964–2974.

39. Reinhart-King CA, Dembo M, Hammer DA (2005) The dynamics and mechanics of endothelial cell spreading. Biophys J 89(1):676–689.

40. Dubin-Thaler BJ, Giannone G, Döbereiner H-G, Sheetz MP (2004) Nanometer analysis of cell spreading on matrix-coated surfaces reveals two distinct cell states and STEPs. Biophys J 86(3):1794–1806.

41. Denisin AK, Pruitt BL (2016) Tuning the Range of Polyacrylamide Gel Stiffness for Mechanobiology Applications. ACS Appl Mater Interfaces 8(34):21893–21902.

42. Choi CK, et al. (2008) Actin and alpha-actinin orchestrate the assembly and maturation of nascent adhesions in a myosin II motor-independent manner. Nat Cell Biol 10(9):1039–1050.

43. Kong F, García AJ, Mould AP, Humphries MJ, Zhu C (2009) Demonstration of catch bonds between an integrin and its ligand. J Cell Biol 185(7):1275–1284.

44. Ponti A, Machacek M, Gupton SL, Waterman-Storer CM, Danuser G (2004) Two distinct actin networks drive the protrusion of migrating cells. Science 305(5691):1782–1786.

45. Rossier O, et al. (2012) Integrins β(1) and β(3) exhibit distinct dynamic nanoscale organizations inside focal adhesions. Nat Cell Biol 14(10):1057–1067.

46. Xu X-P, et al. (2016) Three-Dimensional Structures of Full-Length, Membrane-Embedded Human α(IIb)β(3) Integrin Complexes. Biophys J 110(4):798–809.

47. Burnette DT, et al. (2011) A role for actin arcs in the leading-edge advance of migrating cells. Nat Cell Biol. doi:10.1038/ncb2205.

48. Gailit J, Ruoslahti E (1988) Regulation of the fibronectin receptor affinity by divalent cations. J Biol Chem 263(26):12927–12932.

49. Smith JW, Piotrowicz RS, Mathis D (1994) A mechanism for divalent cation regulation of beta 3-integrins. J Biol Chem 269(2):960–967.

50. Mould AP, Akiyama SK, Humphries MJ (1995) Regulation of integrin alpha 5 beta 1-fibronectin interactions by divalent cations. Evidence for distinct classes of binding sites for Mn2+, Mg2+, and Ca2+. J Biol Chem 270(44):26270–26277.

51. Choquet D, Felsenfeld DP, Sheetz MP (1997) Extracellular matrix rigidity causes strengthening of integrin-cytoskeleton linkages. Cell 88(1):39–48.

52. Suter DM, Errante LD, Belotserkovsky V, Forscher P (1998) The Ig superfamily cell adhesion molecule, apCAM, mediates growth cone steering by substrate-cytoskeletal coupling. J Cell Biol 141(1):227–240.

53. Coyer SR, et al. (2012) Nanopatterning reveals an ECM area threshold for focal adhesion assembly and force transmission that is regulated by integrin activation and cytoskeleton tension. J Cell Sci 125(Pt 21):5110–5123.

54. Sabass B, Gardel M, Waterman CM, Schwarz US (2008) High resolution traction force microscopy based on experimental and computational advances. Biophys J 94(1):207–220.

55. Aratyn-Schaus Y, Oakes PW, Stricker J, Winter SP, Gardel ML (2010) Preparation of compliant matrices for quantifying cellular contraction. J Vis Exp (46):e2173–e2173.

56. Tseng Q, et al. (2011) A new micropatterning method of soft substrates reveals that different tumorigenic signals can promote or reduce cell contraction levels. Lab Chip 11(13):2231–2240.

57. Underhill PT, Doyle PS (2004) On the coarse-graining of polymers into bead-spring chains. Journal of Non-Newtonian Fluid Mechanics 122(1-3):3–31.

58. Hotulainen P, Lappalainen P (2006) Stress fibers are generated by two distinct actin assembly mechanisms in motile cells. J Cell Biol 173(3):383–394.

59. Pereverzev YV, Prezhdo OV, Forero M, Sokurenko EV, Thomas WE (2005) The Two-Pathway Model for the Catch-Slip Transition in Biological Adhesion. Biophys J 89(3):1446–1454.

60. Pereverzev YV, Prezhdo OV, Thomas WE, Sokurenko EV (2005) Distinctive features of the biological catch bond in the jump-ramp force regime predicted by the two-pathway model. Phys Rev E Stat Nonlin Soft Matter Phys 72(1 Pt 1):010903.

61. Li Y, Bhimalapuram P, Dinner AR (2010) Model for how retrograde actin flow regulates adhesion traction stresses. J Phys Condens Matter 22(19):194113.

